# DIFS: Discriminative Feature Selection for Cell Clustering based on Single-Cell RNA Sequencing Data

**DOI:** 10.1101/2025.03.26.645625

**Authors:** Kun Liu, Jinpeng Liu, Chi Wang

## Abstract

A key challenge in single-cell RNA sequencing (scRNA-seq) analysis is clustering cells based on their expression profiles. Effective clustering requires selecting the most informative gene features whose varying expression levels in different cell types can be used to discriminate between different cell types. This study introduces DIFS, a novel statistical framework designed to enhance discriminative feature selection for scRNA-seq-based cell clustering. DIFS operates in two stages. In the first stage, a modified dip test identifies genes with significant multimodal expression patterns, as these are likely to have different expression levels in different cell types. In the second stage, cells are clustered based on the selected features from stage one, and additional cluster-specific features are identified, capturing genes that may be expressed in only one cell cluster. Through real data analysis, we demonstrate that DIFS improves the accuracy of cell type classification and enhances the understanding of cellular heterogeneity in single-cell studies.

## 1 Introduction

Single-cell RNA sequencing (scRNA-seq) has revolutionized the understanding of cellular diversity and gene expression patterns within tissues, offering a high-resolution view of cell-to-cell variability. This technology enables the identification of novel cell types and provides insight into regulatory mechanisms at a previously unattainable level of detail. Cell clustering, which clusters cells based on their expression profiles, is an important task in scRNA-seq data analysis. It provides valuable information for downstream cell type identification, differential expression analysis, and cell trajectory inference.

Gene feature selection is a critical step in cell clustering. As many genes have similar expression levels across different cell types, they do not contribute to cell clustering. Therefore, only including informative gene features that are differentially expressed in different cell types can substantially enhance the signal to noise ratios, leading to more accurate cell clustering results [19]. Various feature selection methods have been proposed for scRNA-seq data. The majority of those methods aim to select genes based on either the mean expression value or the variance, or both. However, genes with high mean expressions are not necessarily differentially expressed in different cell types. For example, house-keeping genes typically have high mean expression values, but are constantly expressed across different cell types. Likewise, genes with large variances may also not contribute to distinguish different cell clusters. We applied the high variability feature selection method in Seurat, a popular software package for scRNA-seq analysis, to the pbmc3k dataset, which is a sample dataset featured on the Seurat website. Gene RBM39 was among the top features identified by Seurat. This gene had a large variance as compared to its mean (Figure 3), which is why it was selected by Seurat. However, Figure 2 left panel shows that it was consistently expressed in almost all cell types, and thus is not very informative to distinguish different cell types. Therefore, mean and variance are only indirectly related to differential expression. Gene features selected based on those criteria may be suboptimal for cell clustering. In addition to methods based on mean and/or variance, there are some other methods that convert this unsupervised feature selection problem into a supervised problem. Those methods first perform an initial clustering to identify preliminary cell clusters and then select features that are differentially expressed between different preliminary cell clusters. A limitation of those methods is that they depend on the accuracy of the preliminary cell clusters, which can be challenging, since the goal of feature selection is to enhance cell clustering.

**Fig. 1.**
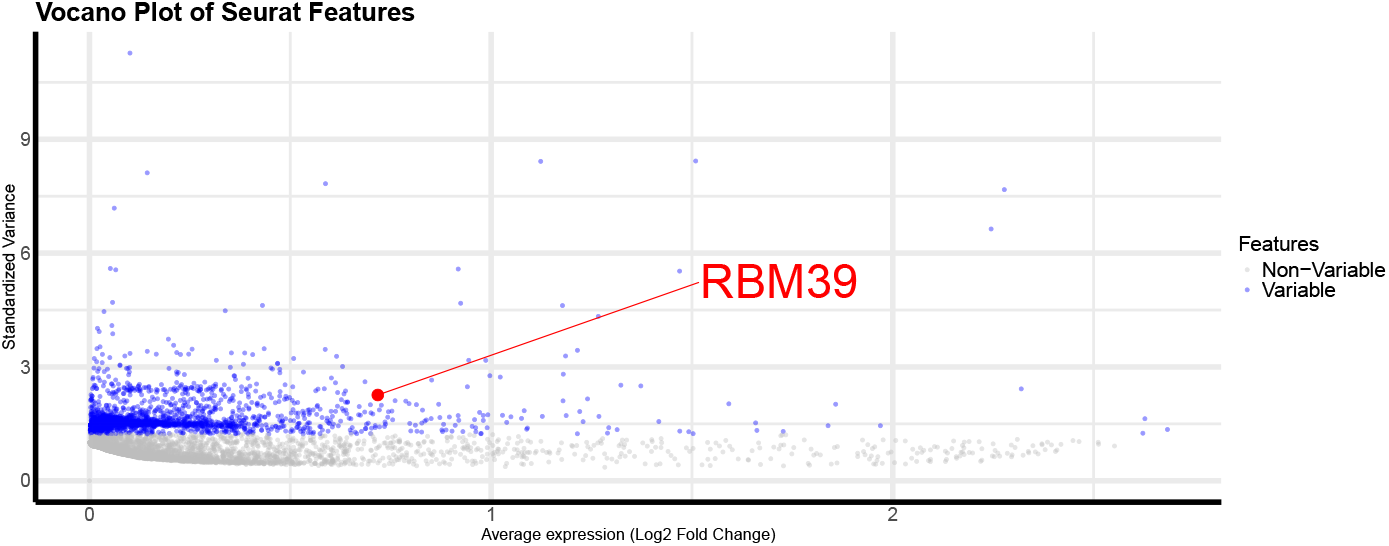
Volcano plot of gene expression variability. Gene ‘RBM39’ is marked as a variable feature by Seurat, but it does not show significant differential expression across cell types.

**Fig. 2.**
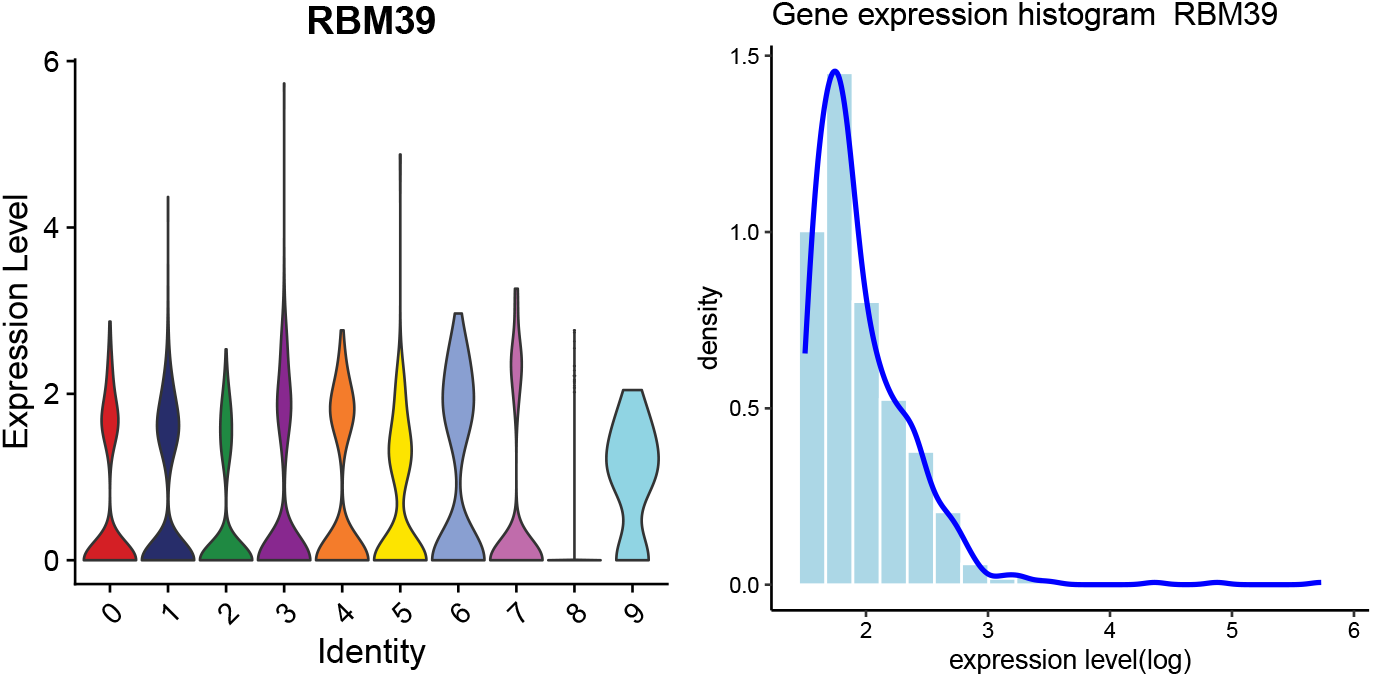
Violin plot and histogram of the feature ‘RBM39’. Despite its high variability, this gene does not provide useful information for distinguishing cell types. The histogram shows the distribution of gene expression levels on a log-transformed scale.

**Fig. 3.**
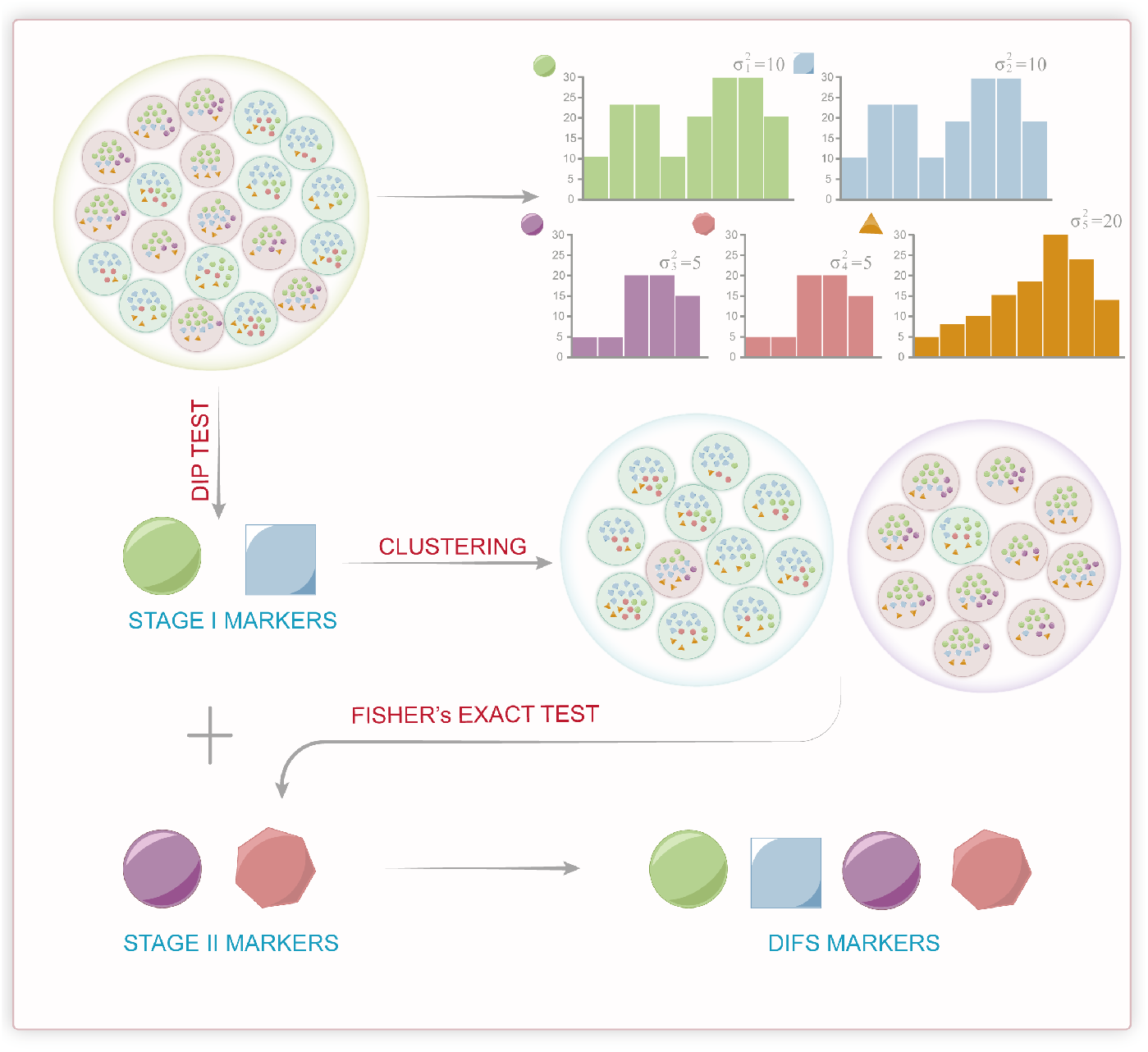
DIFS work flow.

For an informative gene feature, since it is differentially expressed in different cell types, its marginal distribution across all cells is a mixture of two or more distributions, coming from the cell type(s) with or without this gene upregulated. Therefore, the marginal distribution is expected to be multimodal. Based on this observation, we propose to select features by assessing the multimodality in their expression distributions, which is a direct approach since multimodality is the characteristic of informative features. In statistics, one method to assess whether a distribution is unimodal or multimodal is the dip test. The dip test, introduced by Hartigan and Hartigan (1985), is a non-parametric statistical method used to. It specifically measures the “dip,” defined as the maximum difference between the empirical distribution function (EDF) of the observed data and the closest unimodal distribution. In essence, the dip statistic quantifies the extent of deviation from unimodality. A larger-than-expected dip statistic, suggests that the distribution may exhibit multiple modes. Under the standard dip test, the null hypothesis posits that the data follow a unimodal distribution. A significant deviation from this null hypothesis, indicated by a larger-than-expected dip statistic, suggests that the distribution may exhibit multiple modes, challenging the assumption of unimodality. In the original dip test, the null distribution is assumed to be a uniform distribution, which is too conservative (see section 2.2).

In this article, we introduce a new framework, named DIscriminative Feature Selection (DIFS), for identifying informative gene features by assessing multimodal expression patterns. A cornerstone of our framework is a modified dip test. We propose to use an empirical normal distribution instead of uniform distribution as the null hypothesis, which is able to substantially enhance the power of the dip test. Based on the modified version of dip test, we propose a two-stage procedure to identify informative features. The performance of DIFS is evaluated and compared to existing methods based on 12 publicly available scRNA-seq datasets.

## 2 Results

### 2.1 Overview

Our assumption is that if a gene is non-informative, it should have the same distribution across all cell types. Genes with multimodal distributions may contain valuable biological information that benefits cell clustering. Our aim is to identify these genes with mixture distributions. However, there is one type of gene that cannot be neglected—genes that are expressed only in specific cell types. These cell type-specific genes also have a unimodal distribution. In light of this, we divided our markers into two types: genes with multimodal distributions, known as stage I markers (4), and cell type-specific genes, known as stage II markers (5).

### 2.2 Performance Comparison between the Modified and Original Dip Tests

The cornerstone of the Differential Feature Selection (DIFS) framework is the identification of stage I markers—genes that exhibit significant multimodal expression patterns across single cells. This is achieved using a modified version of Hartigan’s Dip Test [7]. In this subsection, we evaluated the performance of the original and modified dip tests through simulations. Specifically, we generated 1000 samples, each consisting of 1000 data points, drawn from a normal distribution with varying means and variances. Since the simulated data are unimodal, this represents a null scenario where p-values from the test are expected to follow a uniform distribution.

We applied the original dip test to the simulated data, and the resulting p-value distribution is shown in Figure 6. The distribution is heavily skewed toward 1, deviating substantially from a uniform distribution, indicating that the original dip test is overly conservative. In practice, this conservatism makes it difficult to identify a sufficient number of stage I markers, as most genes yield p-values of 1, rendering it infeasible to rank them effectively.

**Fig. 4.**
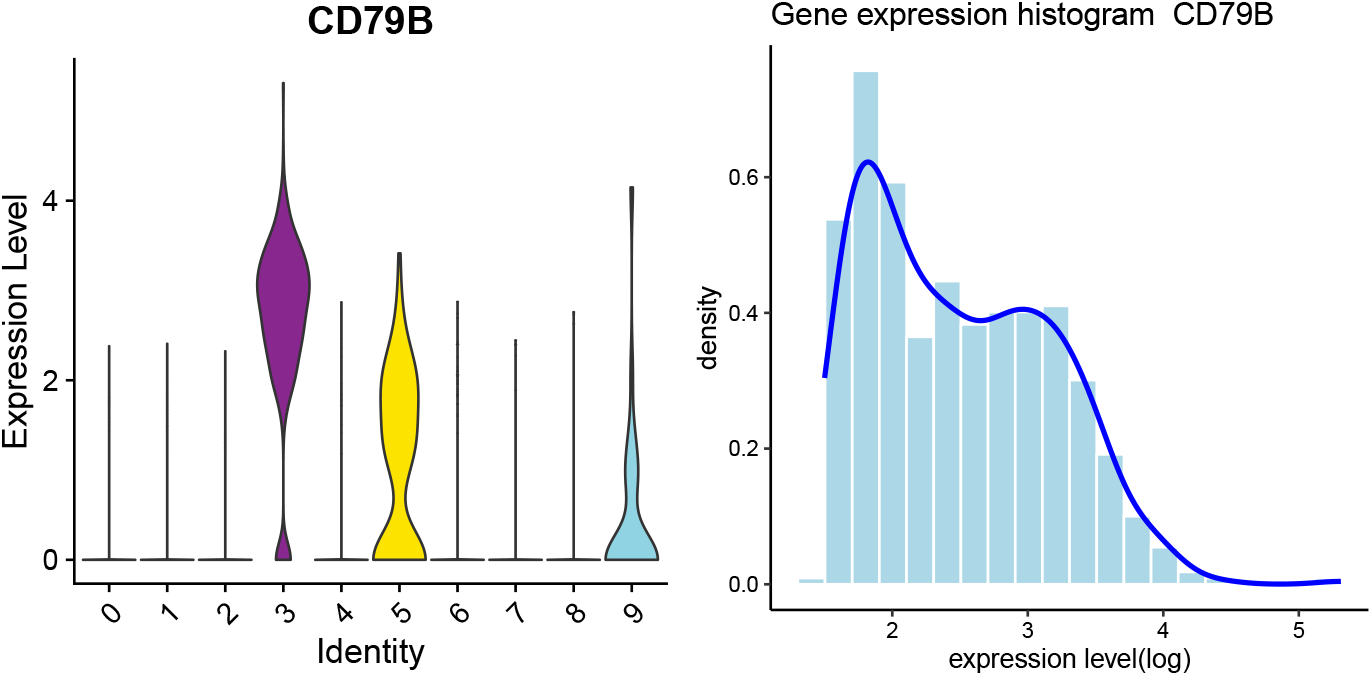
Violin plot and histogram displaying expression levels of stage I marker gene ‘CD79B’ across various clusters. The histogram shows the distribution of gene expression levels on a log-transformed scale, indicating the density and distribution of expression per cell identity.

**Fig. 5.**
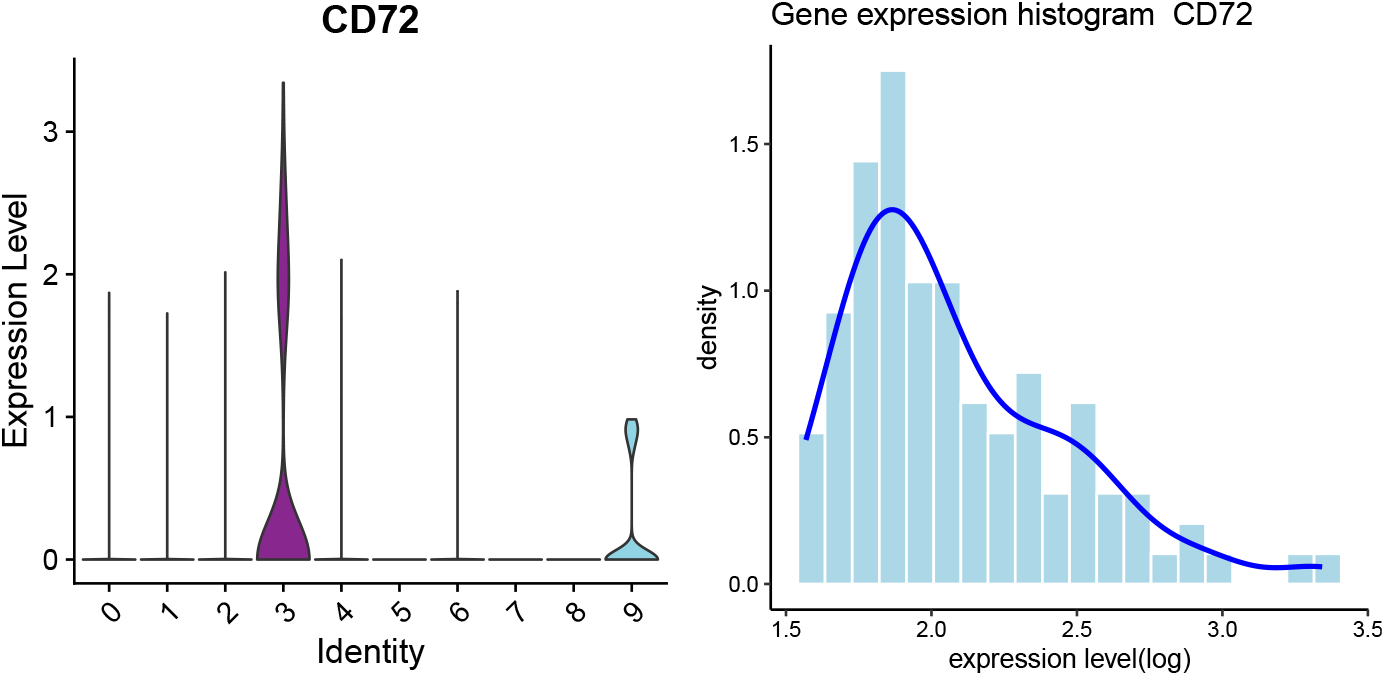
Violin plot and histogram displaying expression levels of stage II marker gene ‘CD72’ across various clusters. The histogram shows the distribution of gene expression levels on a log-transformed scale, highlighting the effective identification of differential markers within cell clusters.

**Fig. 6.**
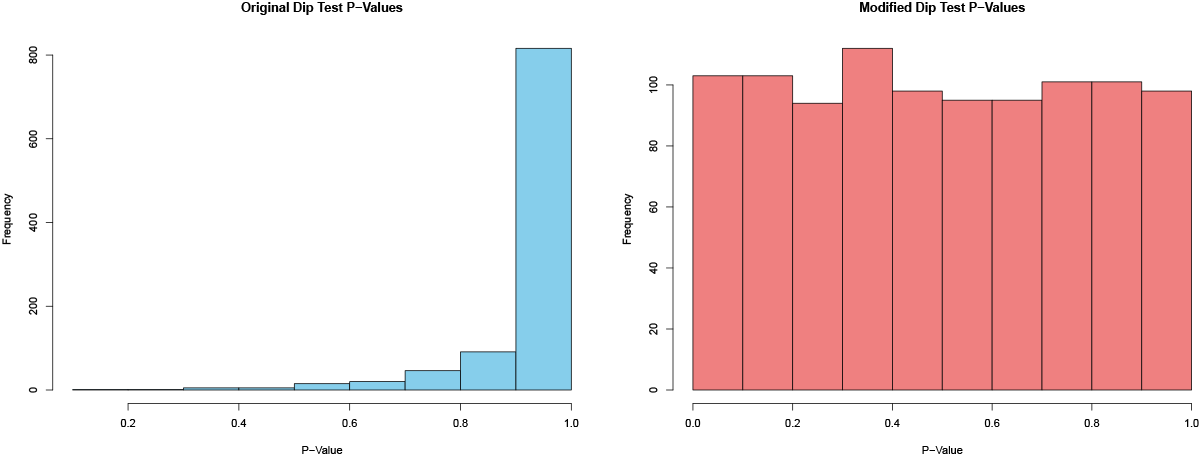
Distribution of Dip Tests’ p-values with uniform underlying distribution

By contrast, the p-values from our modified dip test were more uniformly distributed between 0 and 1 (Figure 6), demonstrating that it is better suited for assessing multimodality. The key difference between the original and modified dip tests lies in how comparison samples are generated when using the R package diptest [14]. The original dip test uses a uniform distribution to generate comparison samples, whereas one assumption of this paper is that gene expression distributions are closer to a normal distribution. Consequently, the modified test generates comparison samples from an empirical normal distribution, which we believe provides a more accurate basis for evaluating multimodality in gene expression data.

### 2.3 Assessing the Performance of DIFS based on Publicly Available Datasets

We applied the proposed marker selection method, DIFS, to 12 publicly available scRNAseq datasets to assess the performance of our method as well as to compare it with existing feature selection methods. Those 12 scRNAseq datasets include both simulated and real datasets with varying sample sizes and complexities, allowing us to comprehensively evaluate the methods performance under various scenarios. The existing feature selection methods under comparison include Seurat, Feast and SC3. To make a fair comparison, we selected 1000 features from each feature selection method. Note that the SC3 method is designed to pick all features except those expressed in either too few or too many cells. To ensure a fair comparison, we randomly selected 1000 features for the SC3 method following the practice in FEAST paper [19].

Gene features identified by a feature selection method were passed to a clustering method for cell clustering based on the selected features. The performances of different feature selection methods were then compared based on the accuracy in cell clustering. For a robust comparison, we considered five clustering methods, including Monocle[16], TSCAN[11], Seurat (both refined Louvain and SLM algorithms were considered), and SC3[12]. In total, twenty different feature selection-clustering method combinations were considered. The performance of each method combination was evaluated based on ARI, FM, Jaccard Index, and Purity, providing a comprehensive analysis of clustering quality.

The ARI values from different method combinations across the 12 datasets are presented in a heatmap Figure 7. Each row of the heatmap corresponds to a dataset, with columns representing different clustering methods such as ‘monocle’, ‘TSCAN’, ‘Refined Louvain’, ‘SLM’, and ‘SC3’. The color intensity in the heatmap transitions from light to dark, corresponding to performance levels from lower to higher ARI values. The feature selection-clustering method combinations are sorted by the average ARI across datasets. Among different feature selection methods, DIFS had the highest or close to the highest ARI values in all 12 datasets. This observation holds for all the clustering methods we considered. Among different clustering methods, the SC3 clustering method tended to provide the highest ARI value for a given feature selection method.

**Fig. 7.**
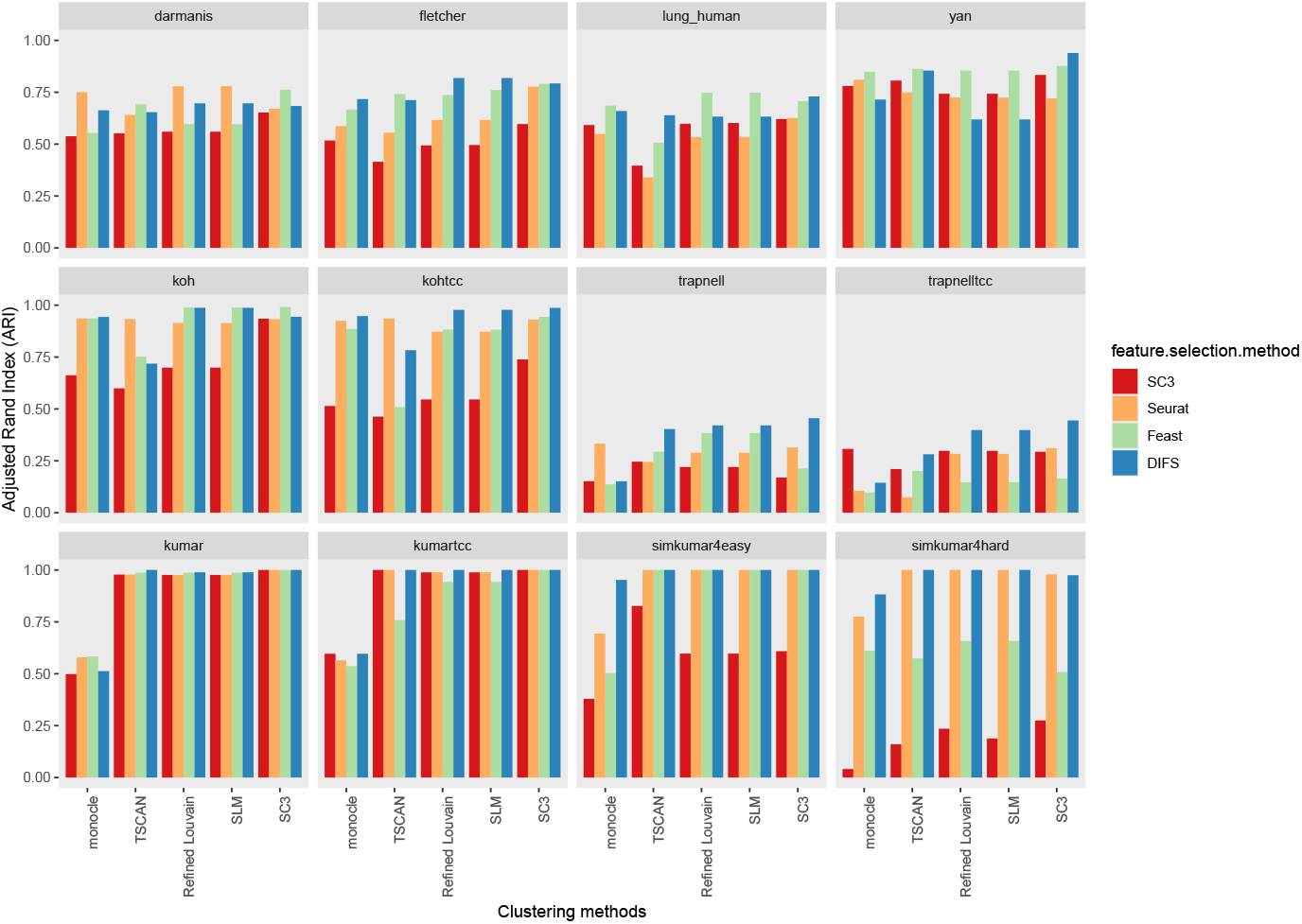
Comparative analysis of clustering performance across various scRNA-seq datasets. Each panel of bar plots corresponds to a different dataset tested with various feature selection and clustering strategies, illustrating the Adjusted Rand Index (ARI) scores achieved.

We also examined datasets where our method demonstrates clearly superior performance compared to other feature selection approaches—for example, the Trapnell dataset. This dataset includes cells at various developmental stages, resulting in subtle gene expression differences across stages. Despite the inherent difficulty of this scenario, the DIFS method effectively identified informative genes and produced more reliable results. In contrast, Seurat may overlook features with low variance, potentially missing important signals. For FEAST, its feature selection step depends on pseudo-clustering prior to performing the F-test. Given the challenging nature of clustering in this dataset, the performance of FEAST is significantly compromised.

To complement the heatmap, a line chart (Figure 8) was generated to present the average ARI across all datasets and all clustering methods for each feature selection method. DIFS has the highest average ARI value. In addition to ARI, Figure 8 also compares the average FM, Jaccard, and purity values across different feature selection methods. The results are consistent, with DIFS having the highest values in all those measures. Therefore, DIFS overall outperformed existing feature selection methods.

**Fig. 8.**
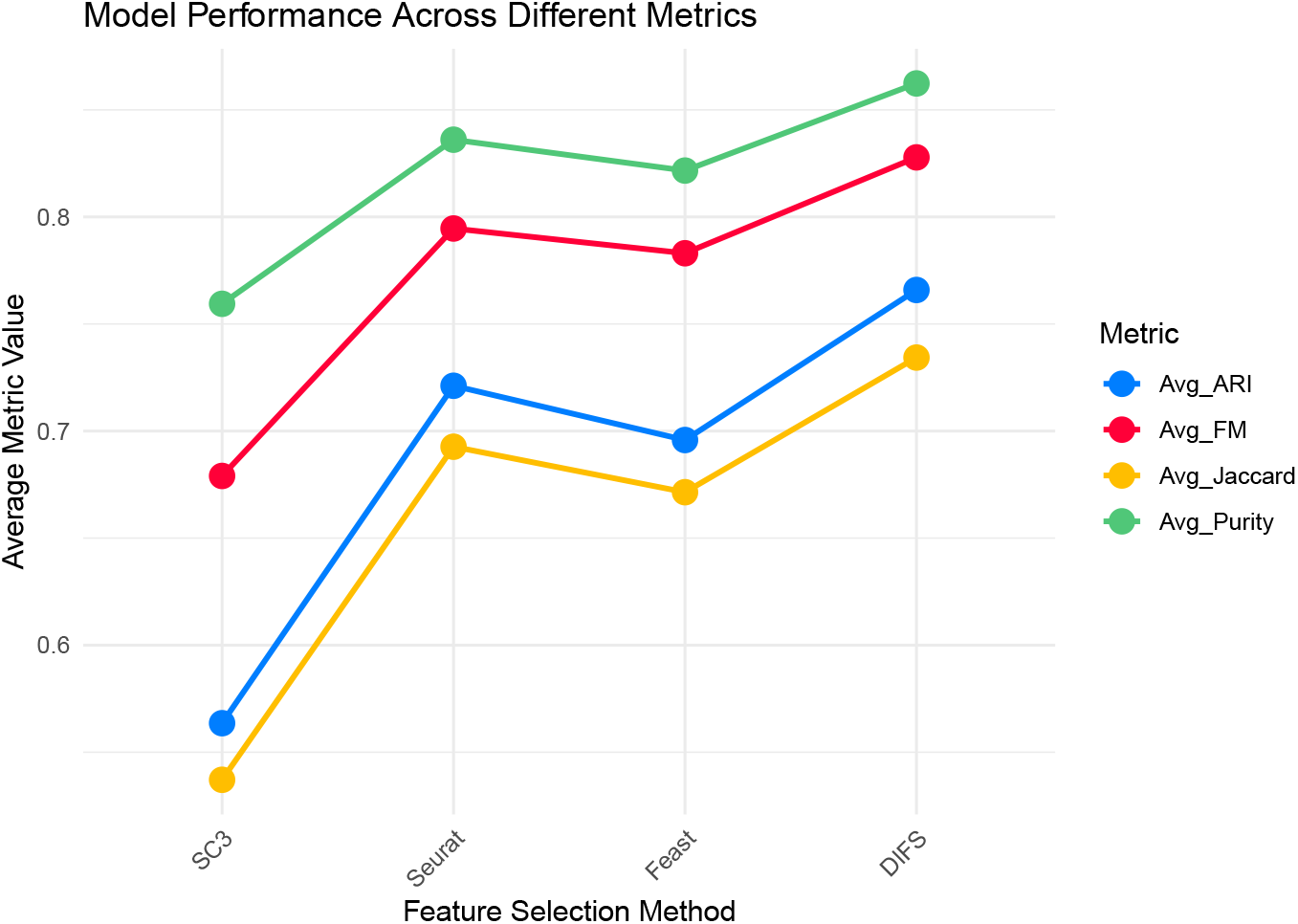

## 3 Methods

### 3.1 A Modified Dip Test

#### 3.1.1 The Original Dip Test

The dip statistic is a measure used to quantify the extent to which a given empirical distribution deviates from being unimodal. To define this rigorously, consider an empirical distribution function *F* (*x*) derived from a sample of data. A distribution function *F* is considered unimodal with a mode *m* if it satisfies two key properties:

- *F* is *convex* on the interval (−∞, *m*], meaning that for any points *x*_1_ < *x*_2_ < *m*, the line segment connecting *F* (*x*_1_) and *F* (*x*_2_) lies above the graph of *F*.
- *F* is *concave* on the interval [*m*, ∞), meaning that for any points *m* < *x*_1_ < *x*_2_, the line segment connecting *F* (*x*_1_) and *F* (*x*_2_) lies below the graph of *F*.

The mode *m* is the point where this transition from convexity to concavity occurs. While the mode is typically unique, it is possible for a distribution to have multiple modes, each corresponding to a different interval where the distribution is unimodal.

To formally define the dip statistic, we first introduce the concept of a distance measure between two distribution functions *F* and *G*, denoted as *p*(*F, G*), which is the maximum absolute difference between them over all possible values of *x*:

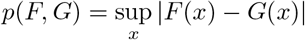

This measure captures the greatest vertical distance between the cumulative distribution functions (CDFs) of *F* and *G* at any point *x*.

Next, consider the class 𝒰 of all unimodal distribution functions, which encompasses all distributions that satisfy the convexity and concavity conditions described above. The dip statistic *D*(*F*) of the empirical distribution function *F* is then defined as the smallest distance between *F* and any unimodal distribution function *G* within the class 𝒰 :

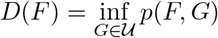

This infimum identifies the closest unimodal distribution to *F* and measures how far *F* deviates from this optimal unimodal fit. The dip statistic *D*(*F*) provides a precise measure of the departure from unimodality:

- If *D*(*F*) = 0, then *F* itself is unimodal, as it lies within the class 𝒰.
- If *D*(*F*) > 0, then *F* is not unimodal, with the value of *D*(*F*) quantifying the extent of this deviation.

In other words, the dip statistic identifies the degree to which the empirical distribution function *F* fails to conform to a unimodal shape. This makes the dip an essential tool for distinguishing between unimodal and multimodal distributions, particularly in complex datasets where such distinctions are crucial for further analysis.

In practice, the dip statistic is used to detect the presence of multiple modes in a distribution, with higher values of the dip indicating a greater likelihood of multimodality. This is particularly useful in applications such as gene expression analysis, where identifying whether a gene’s expression follows a single pattern or multiple distinct patterns is critical for understanding underlying biological processes.

#### 3.1.2 The Modification

While the traditional Dip Test employs a uniform distribution as the null hypothesis, this choice may not be ideal in all contexts—particularly in gene expression studies. When analyzing gene expression data, we assume that if a gene expresses consistently across all cell types without differential expression, the distribution of its expression levels should follow a unimodal Gaussian distribution. In this scenario, the use of a uniform distribution as the null hypothesis fails to capture this underlying biological assumption, making it less suitable for detecting deviations from unimodality.

To address this, we modify the Dip Test by adopting an empirical normal distribution as the null hypothesis. This approach aligns more closely with the biological expectation that a gene uniformly expressed across cell types should exhibit a unimodal Gaussian distribution. By comparing the empirical distribution of gene expression data against this empirical normal distribution, we can more accurately detect deviations from the expected unimodal pattern, thereby enhancing our ability to identify genes with differential expression or other biologically significant patterns.

#### 3.1.3 Advantages of Using an Empirical Normal Distribution

The adoption of an empirical normal distribution as the null hypothesis for the Dip Test offers several key advantages in the context of gene expression analysis:

- **Increased Test Power:** One of the most significant benefits is the increased sensitivity of the Dip Test. By aligning the null hypothesis with the true underlying distribution, the test gains greater power to detect deviations from unimodality. This enhanced sensitivity allows us to more effectively identify genes that deviate from a unimodal distribution, potentially revealing important differential expression patterns.
- **Identification of Invariant Genes:** In practice, some genes may be invariant in certain cell types but not across all types. These genes often exhibit distributions with many cells showing low expression, leading to a log-normal-like distribution. By using an empirical normal distribution as the null hypothesis, the Dip Test becomes more adept at detecting such cases, yielding a higher dip statistic for genes that deviate from the expected unimodal Gaussian distribution.
- **Addressing Log-Normal Distributions:** Gene expression data frequently display log-normal distributions, particularly when there is a mix of high and low expression levels across cells. Despite applying a log transformation to the data, which is a common practice to normalize expression levels, we often observe that the resulting distribution still exhibits a log-normal pattern. This residual log-normal behavior suggests that the data retain certain characteristics that deviate from simple unimodality. The empirical normal distribution helps to better identify these situations, where the assumption of unimodality under a uniform distribution might not hold. By adapting the null distribution to an empirical normal, the Dip Test can more effectively distinguish between truly unimodal distributions and those with subtle deviations due to log-normal behavior, even after log transformation.

### 3.2 A Two-Stage Feature Selection Procedure

#### 3.2.1 Stage I Marker Selection based on the Modified Dip Test

Our modifications make the Dip Test a more powerful tool for scRNA-seq data analysis. Genes exhibiting low p-values from this refined approach are identified as potential stage I markers. These markers are crucial as they reflect pronounced multimodal expression patterns indicative of diverse cellular states, playing a key role in subsequent clustering and analysis.

We adopt a nuanced selection process, ranking potential stage I markers by their p-values and selecting a predefined number (e.g., top 500, 1000, or 2000) for further analysis. This method accounts for the overall distribution of p-values and includes genes with moderately higher p-values, which may still hold significant biological insights.

Acknowledging the challenge in determining the optimal number of stage I markers to retain, we introduce an auxiliary decision-making tool. This tool allows users to specify potential gene set sizes, performs clustering for each size, and constructs a linear model correlating clustering labels with gene expression levels. It then calculates the mean squared error (MSE) for each gene set size, recommending the set with the minimal MSE for use in clustering.([19])

#### 3.2.2 Stage II Marker Selection based on the Fisher’s Exact Test

After deciding the optimal set of stage I markers, we cluster the data with the selected stage I markers using popular clustering methods such as Seurat [18] and SC3 [12]. We then apply Fisher’s exact test to each gene to find cluster-specific markers. Instead of converting all non-zero counts to ‘1’, we binarize the data into high expression and low expression with a cutoff of ln(5) on the log-transformed data. For each gene, we perform the test on the cluster where its expression is highest against other clusters. We also considered performing Fisher’s test on the highest expressed cluster against the second highest, but this approach could miss valuable genes. This is because the number of clusters used in our experiments may not reflect the true biological subtypes. There might be more subtypes that are not of interest to researchers or closely related clusters where a gene could be a marker for a cell type but is spread across subtypes. In such cases, comparing only the top expressed clusters would neglect important markers. After the Fisher’s exact test, we rank the genes based on p-values and mix stage I and stage II markers using a ratio from 1:3 to 3:1. For example, for a ratio of 1:3, the gene list is arranged as one stage I marker followed by three stage II markers, then another stage I marker followed by three stage II markers. Since we focus on the top 500, 1000, or 2000 features, we have sufficient stage I and stage II markers to mix them in different ratios and provide the user with the optimal gene set as the selected markers.

### 3.3 Datasets

We evaluated 12 publicly available scRNA-seq datasets, including Darmins, Fletcher, Lung Human, Yan, Koh, KohTCC, Trapnell, TrapnellTCC, Kumar, KumarTCC, SimKumar4easy, and SimKumar4hard. The first four datasets—Darmins, Fletcher, Lung Human, and Yan—were also used in the FEAST paper [19], while the remaining eight were introduced in another study [5]. Notably, SimKumar4easy and SimKumar4hard are simulated datasets based on real data. In particular, SimKumar4hard is designed to reflect challenging conditions by using the Splatter package [22], simulating a complex environment with four subpopulations and varying probabilities of differential expression. For all datasets, transcript compatibility counts (TCCs) were also estimated and used as an alternative to the gene-level count matrix for input into the clustering algorithms.

### 3.4 Data Processing

The input data is structured as a gene expression matrix, where rows represent genes, columns represent cells, and each entry corresponds to the expression level of a particular gene in a specific cell. To ensure reliable analysis, we filter the gene expression dataset to include only genes expressed in a substantial number of cells above a threshold of log 5. We chose the cutoff based on our observation across different datasets. This preprocessing step reduces noise and enhances the reliability of the Dip Test.

### 3.5 Evaluation Metrics

We employed four metrics, including Adjusted Rand Index (ARI) [9], Fowlkes-Mallows (FM) Index [1], Jaccard index [10], and Purity [6], to provide a comprehensive evaluation of clustering performance. The definitions of the metrics are as follows.

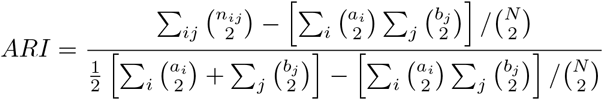

where *n*_*ij*_ is the number of objects in both clusters *i* and *j, a*_*i*_ and *b*_*j*_ are the objects in clusters *i* and *j*, respectively, and *N* is the total number of objects. The ARI ranges from -1 (no agreement) to 1 (perfect agreement), with 0 indicating random chance.

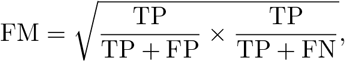

where TP is the number of pairs of points in the same cluster in both clusterings, FP is the number of pairs of points in the same cluster in one clustering but not the other, and FN is the number of pairs of points in different clusters in one clustering but in the same cluster in the other.

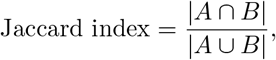

where *A* and *B* are two sets, |*A* ∩ *B*| is the number of elements in both sets, and |*A* ∪ *B*| is the number of elements in either set.

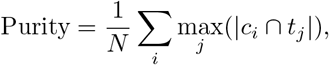

where *N* is the total number of elements, *c*_*i*_ is the set of elements in cluster *i, t*_*j*_ is the set of elements in class *j*, and |*c*_*i*_ ∩ *t*_*j*_| is the number of elements of class *j* in cluster *i*.

## 4 Discussion

This study presented a comprehensive comparison of clustering algorithms across a range of datasets, with a particular focus on the performance of the DIFS method. Our findings, as illustrated in the heatmaps and the line chart, unequivocally demonstrate the DIFS method’s superior clustering performance, evidenced by consistently higher ARI scores.

The DIFS method’s robustness across various datasets, including ‘kumar’, ‘trapnell’, and ‘koh’, highlights its adaptability and reliability in deciphering complex biological data. The darker color intensity in the heatmap sections corresponding to the DIFS method not only signifies its effectiveness but also suggests its potential in uncovering subtle patterns within the data that other methods may overlook.

Furthermore, the analysis revealed that the combination of feature selection strategies with the DIFS method plays a pivotal role in enhancing clustering accuracy. This underscores the importance of methodological synergy in achieving optimal results. The significant performance leap facilitated by the DIFS method could pave the way for novel biological insights, enhancing our understanding of underlying genetic mechanisms.

Future research should explore the integration of the DIFS method with other data analysis frameworks, potentially extending its applicability beyond clustering to other areas of computational biology. Moreover, investigating the method’s performance on even larger and more heterogeneous datasets could further validate its scalability and effectiveness.

In conclusion, the DIFS method represents a significant advancement in the clustering of scRNA-seq data, offering a powerful tool for researchers in the quest to unravel the complexities of genomic data.

## References

[1] Tim Anderson and Travis J. Wheeler. An optimized fm-index library for nucleotide and amino acid search. Algorithms for Molecular Biology, 16(25), 2021.

[2] Jiahua Chen and Pengfei Li. Hypothesis test for normal mixture models: The em approach. 2009.

[3] Jiahua Chen and Pengfei Li. Hypothesis test for normal mixture models: The em approach. The Annals of Statistics, 37(6A):2523–2542, 2009.

[4] Rui Duan, Yang Ning, Shuang Wang, Bruce G Lindsay, Raymond J Carroll, and Yong Chen. A fast score test for generalized mixture models. Biometrics, 76(3):811–820, 2020.

[5] Angelo Duó, Mark D. Robinson, and Charlotte Soneson. A systematic performance evaluation of clustering methods for single-cell rna-seq data [version 3; peer review: 2 approved]. F1000Research, 7:1141, 2018.

[6] Paolo Ferragina and Giovanni Manzini. Opportunistic data structures with applications. In Proceedings of the 41st Annual Symposium on Foundations of Computer Science, pages 390–398, 2000.

[7] John A Hartigan and Pamela M Hartigan. The dip test of unimodality. The annals of Statistics, pages 70–84, 1985.

[8] Jian Hu, Xiangjie Li, Gang Hu, Yafei Lyu, Katalin Susztak, and Mingyao Li. Iterative transfer learning with neural network for clustering and cell type classification in single-cell rna-seq analysis. Nature machine intelligence, 2(10):607–618, 2020.

[9] Vandra L Huber. Effects of task difficulty, goal setting, and strategy on performance of a heuristic task. Journal of Applied Psychology, 70(3):492, 1985.

[10] Paul Jaccard. Étude comparative de la distribution florale dans une portion des alpes et des jura. Bulletin de la Société Vaudoise des Sciences Naturelles, 37:547–579, 1901.

[11] Zhicheng Ji and Hongkai Ji. Tscan: Pseudo-time reconstruction and evaluation in single-cell rna-seq analysis. Nucleic Acids Research, 44(13):e117, 2016.

[12] Vladimir Kiselev, Kirill Kirschner, Michael T. Schaub, Theresa Andrews, Andrew Yiu, T. Alan Righelli, and Kerstin Hemberg. Sc3: consensus clustering of single-cell rna-seq data. Nature Methods, 14:483–486, 2017.

[13] Wenjing Ma, Kenong Su, and Hao Wu. Evaluation of some aspects in supervised cell type identification for single-cell rna-seq: classifier, feature selection, and reference construction. Genome biology, 22:1–23, 2021.

[14] Martin Maechler. diptest: Hartigan’s Dip Test Statistic for Unimodality - Corrected, 2023. R package version 0.77-0.

[15] Paul D McNicholas. Mixture model-based classification. Chapman and Hall/CRC, 2016.

[16] Xiaojie Qiu, Qiaolin Mao, Ying Tang, Lauren Wang, Rachael Chawla, Hector Pliner, Jonathan Y. Trapnell, and Cole Trapnell. Reversed graph embedding resolves complex single-cell trajectories. Nature Methods, 14:979–982, 2017.

[17] Charlotte Soneson and Mark D Robinson. Bias, robustness and scalability in single-cell differential expression analysis. Nature methods, 15(4):255–261, 2018.

[18] Tim Stuart, Andrew Butler, Paul Hoffman, Christoph Hafemeister, Efthymia Papalexi, William M. Mauck, Yuhan Hao, Marlon Stoeckius, Peter Smibert, and Rahul Satija. Comprehensive integration of single-cell data. Cell, 177(7):1888– 1902.e21, 2019.

[19] Kenong Su, Tianwei Yu, and Hao Wu. Accurate feature selection improves single-cell rna-seq cell clustering. Briefings in bioinformatics, 22(5):bbab034, 2021.

[20] Cole Trapnell, Davide Cacchiarelli, Jonna Grimsby, Prapti Pokharel, Shuqiang Li, Michael Morse, Niall J Lennon, Kenneth J Livak, Tarjei S Mikkelsen, and John L Rinn. Pseudo-temporal ordering of individual cells reveals dynamics and regulators of cell fate decisions. Nature biotechnology, 32(4):381, 2014.

[21] Sheng Wang, Angela Oliveira Pisco, Aaron McGeever, Maria Brbic, Marinka Zitnik, Spyros Darmanis, Jure Leskovec, Jim Karkanias, and Russ B Altman. Leveraging the cell ontology to classify unseen cell types. Nature communications, 12(1):5556, 2021.

[22] Luke Zappia, Belinda Phipson, and Alicia Oshlack. Splatter: simulation of single-cell rna sequencing data. Genome Biology, 18(1):174, 2017.

[23] Ying Zhao and George Karypis. Criterion functions for document clustering: Experiments and analysis. Machine Learning, 55(3):311–331, 2001.

